# Edible Bird’s Nest Enhances Memory and Learning in a Rat Model of Sporadic Alzheimer’s Disease

**DOI:** 10.1101/2025.05.22.655527

**Authors:** Nurul Husna Ibrahim, Mohamad Fairuz Yahaya, Seong Lin Teoh, Wael Mohamed, Chua Kien Hui, Azizah Ugusman, Jaya Kumar

## Abstract

Alzheimer’s disease (AD) has emerged as a major global health concern, with the glutamate hyperexcitability hypothesis suggesting that synaptic dysfunction and neuronal death are key contributors to cognitive impairments. Despite decades of research, current AD treatments only manage symptoms without offering a cure. This study investigates the potential of edible bird’s nest (EBN) as a therapeutic intervention for mitigating cognitive decline in a streptozotocin (STZ)-induced model of sporadic AD. Male Sprague-Dawley rats were randomly assigned to seven groups: control, sham, STZ, low EBN (LEBN, 300 mg/kg), medium EBN (MEBN, 600 mg/kg), high EBN (HEBN, 1200 mg/kg), and memantine (MEM, 10 mg/kg). On day 1, rats received a 3 mg/kg intracerebroventricular (ICV) injection of STZ and were allowed a 7-day recovery period. Following recovery, the rats were treated with their respective interventions daily via oral gavage for 21 days. Behavioral performance was assessed using the Open Field (OF), Novel Object Recognition Test (NORT), and Morris Water Maze (MWM). After behavioral testing, animals were euthanized, and hippocampal tissues were processed for Congo red staining, Bielschowsky staining, and immunohistochemistry to evaluate the expression of NR1, NR2A, NR2B, Glial fibrillary acidic protein (GFAP), and excitatory amino acid transporter 2 (EAAT2). STZ-treated rats exhibited significant declines in recognition memory, spatial learning, and increased anxiety in the open field, along with elevated Congo red and Bielschowsky staining, and higher expression of N-Methyl-D-aspartate receptor (NMDAR) subunits (NR1, NR2A, NR2B), GFAP, and EAAT2, indicative of astrogliosis and glutamate excitotoxicity. Treatment with MEBN (600 mg/kg) significantly improved exploratory behavior, memory, and cognition in the ICV-STZ rats. MEBN effectively attenuated cognitive decline and modulated hippocampal markers, yielding results comparable to the positive control, memantine. These findings underscore the potential of EBN as a promising therapeutic approach for AD and highlight the increasing relevance of traditional medicine in the treatment of neurodegenerative diseases.

## Introduction

In 2022, more than 55 million people have dementia worldwide, which sums up nearly 10 million new cases yearly and Alzheimer’s disease (AD) contributes to more than 80% of these dementia cases by virtue of the indistinguishable advancement of AD treatments [1]. In addition, dementia ranked as the 5th leading cause of global death in 2016 as the death rate caused by dementia accounted for 2.4 million deaths [2]. From year 2000 to 2017, the deaths caused by heart disease have decreased by 9% while the deaths caused by AD increased by 145% [3]. In Asia, the escalating prevalence of AD is evident, with East Asia currently having the highest estimated number of Aβ-positive AD dementia cases, followed by South Asia, indicating a growing healthcare and caregiving burden across the continent [4].

Sporadic AD (SAD) is the most common form of AD, accounting for about 90-95% of all cases. It is amongst the most complex neurodegenerative diseases, signed by amyloid-β plaques and neurofibrillary tangles which result in cognitive and memory loss [5]. Unlike familial AD (FAD), the aetiologies of SAD are not inculpated by an individual gene or pathway but rather constitute an abstruse network of causatives, including glutamate excitotoxicity, neuroinflammation, oxidative stress, mitochondrial dysfunction, and cholinergic disruptions [6]. To model these complex pathological features of SAD in preclinical research, various experimental approaches have been developed. Among these, the use of streptozotocin (STZ) has gained significant traction. STZ is a glucosamine-nitrosourea compound widely used to induce experimental diabetes and neurodegeneration in animal models. Its toxicity toward insulin-producing pancreatic beta cells has made it a valuable tool for diabetes research, while its intracerebroventricular (ICV) administration in rodents serves as a model for SAD, by disrupting brain insulin signaling and promoting oxidative stress and neuroinflammation [7].

Currently, there is no definitive cure for SAD, treatment primarily aims to manage symptoms and slow the progression of the disease. Research has explored various strategies including inhibition of enzymes that lower amyloid levels, tau tangles and neuroinflammation in the brain [8]. Medications like cholinesterase inhibitors (donepezil, rivastigmine) and glutamate regulators (memantine) are used to temporarily improve or stabilize cognitive function (Fig 1). It has been identified oxidative stress and cerebrovascular dysfunction as key contributors to the pathogenesis of SAD and therapy targets [9].

**Fig 1.**
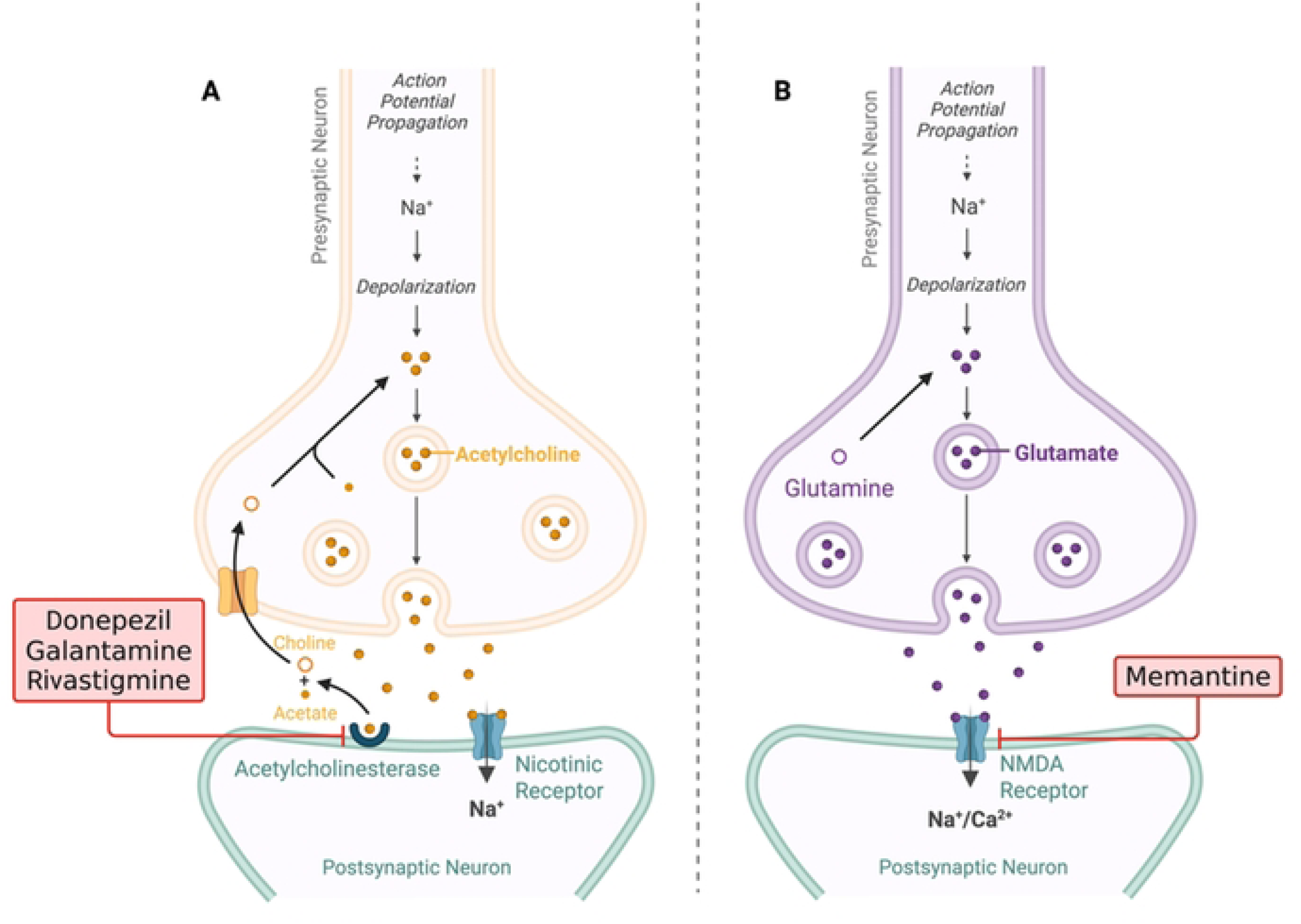
Alzheimer’s Disease - Current Treatments. Generated by BioRender.

Glutamate pathway involving NMDAR is one of the key factors in synaptic plasticity critical for learning and memory development. However, in SAD, this pathway seems to be impaired due to the overstimulation of NMDARs at the glutamatergic tripartite synapse, perturbing the normal glutamate release, uptake and receptors’ activation [10]. The glutamate hypothesis of SAD suggests that abnormalities in the glutamate system may play a crucial role in SAD pathophysiology and could be a potential target for clinical treatments [11]. Research has shown that reduced glutamate neurotransmission is associated with impaired learning and memory in SAD patients [11]. Furthermore, failure to regulate glutamate intake has been implicated in neurological diseases such as SAD [12]. Additionally, recent findings have highlighted glutamate-induced excitotoxicity as a significant breakthrough in understanding the pathogenesis of various neurodegenerative diseases, including SAD [13].

After extensive evaluation of the AD treatment development pipeline, alternative interventions, particularly natural products, have been widely explored in both preclinical and clinical trials. Natural polyphenolic phytochemicals, owing to their potential neuroprotective properties and reduced adverse effect profiles compared to synthetic drugs, present a promising avenue for targeting multiple pathological pathways, particularly those involved in neuroinflammation and oxidative stress. Among these promising natural products, edible bird’s nest (EBN) demonstrates potential neuroprotective properties, attributed in part to its high sialic acid content [14], and is recognized as a traditional tonic food with pleiotropic benefits, including antioxidant and anti-aging effects, which may mitigate oxidative stress, and neuroinflammation in neurodegenerative diseases [15].

This study aimed to evaluate the neuroprotective potential of EBN in a STZ-induced rat model of AD-like phenotypes. We investigated the effects of distinct doses of EBN (low, mid, high) on key AD-related pathologies, including cognitive impairment, amyloid beta plaque formation, neurotangle formation, and glutamate hyperexcitability in the hippocampus. To provide a comparative analysis, we also included a memantine-treated group as a positive control. Memantine, an N-methyl-D-aspartate (NMDA) receptor antagonist, is used to treat moderate to severe AD by regulating the activity of glutamate, a neurotransmitter that is overactive in AD and contributes to neuronal damage. Utilizing a focused experimental design, we sought to determine the most effective EBN dose in mitigating these AD-like phenotypes, and to assess its efficacy relative to memantine.

## Methods

### Animal Preparation

A total of twenty-one (sample size calculated based on Serdar et al. 2021) adult male (age of 12-13 weeks) Sprague-Dawley rats (250-300 g) were obtained from the Laboratory Animal Research Unit (LARU), Universiti Kebangsaan Malaysia (UKM) and housed in Animal Laboratory in separate cages with controlled environment such as temperatures between 25-27°C and 12 hours day and night cycles. The rats were fed with standard rat pellets and tap water ad libitum. Rats were acclimatised for at least 7 days in the animal laboratory before undergoing any procedure. All procedures were carried out in accordance with the institutional guidelines for animal research surgical procedures of the Universiti Kebangsaan Malaysia Research and Animal Ethics Committee (UKMAEC) bearing approval number (FISIO/FP/2020/JAYAKUMAR/16-JAN./1075-JAN.-2020-JAN.-2022).

Twenty-one rats were randomly assigned into seven groups (n=3 per group) following a single bilateral intracerebroventricular (ICV) injection of streptozotocin (STZ, 3 mg/kg) as per Retinasamy et al. [16]. The groups were: Control (no procedure, treated with 0.9% NaCl), Sham (ICV-artificial cerebrospinal fluid (aCSF), treated with 0.9% NaCl), STZ (ICV-STZ, treated with 0.9% NaCl), LEBN (ICV-STZ, treated with 300 mg/kg edible bird’s nest [EBN]), MEBN (ICV-STZ, treated with 600 mg/kg EBN), HEBN (ICV-STZ, treated with 1200 mg/kg EBN), and MEM (ICV-STZ, treated with 10 mg/kg memantine as positive control; Bahramian et al., 2016). All treatments were administered once daily via oral gavage, starting 7 days post-ICV to allow for recovery, and continued until the probe trial day of the Morris Water Maze (MWM), the final day of the study.

### Intracerebroventricular Microinjection of Streptozotocin

For ICV-STZ administration, rats were anesthetized with intraperitoneal ketamine (60 mg/kg) and xylazine (10 mg/kg) on Day 1, following a 7-day habituation period [17] (Fig 2). Body weight was recorded weekly throughout the study. Using a stereotaxic apparatus, each rat was secured with the head flat and stabilized. Ophthalmic ointment was applied to prevent corneal drying, and the body was placed on a thermal pad. After shaving and disinfecting the scalp, a midline incision was made to expose the skull. The bregma was located and used as a reference for injection coordinates: 0.9 mm posterior, ±1.5 mm lateral, and 3.6 mm deep [16]. Small burr holes were drilled, and 5 µl of either STZ (3 mg/kg) or aCSF was bilaterally injected using a 33-gauge needle attached to a Hamilton syringe at a flow rate of 500 nl/min [18]. The syringe was left in place for 5 minutes post-injection to minimize reflux. The incision was closed with surgical staples, antiseptic was applied, and rats received diluted Uphamol for 3 days. A 7-day recovery period was allowed before initiating behavioral testing.

**Fig 2.**
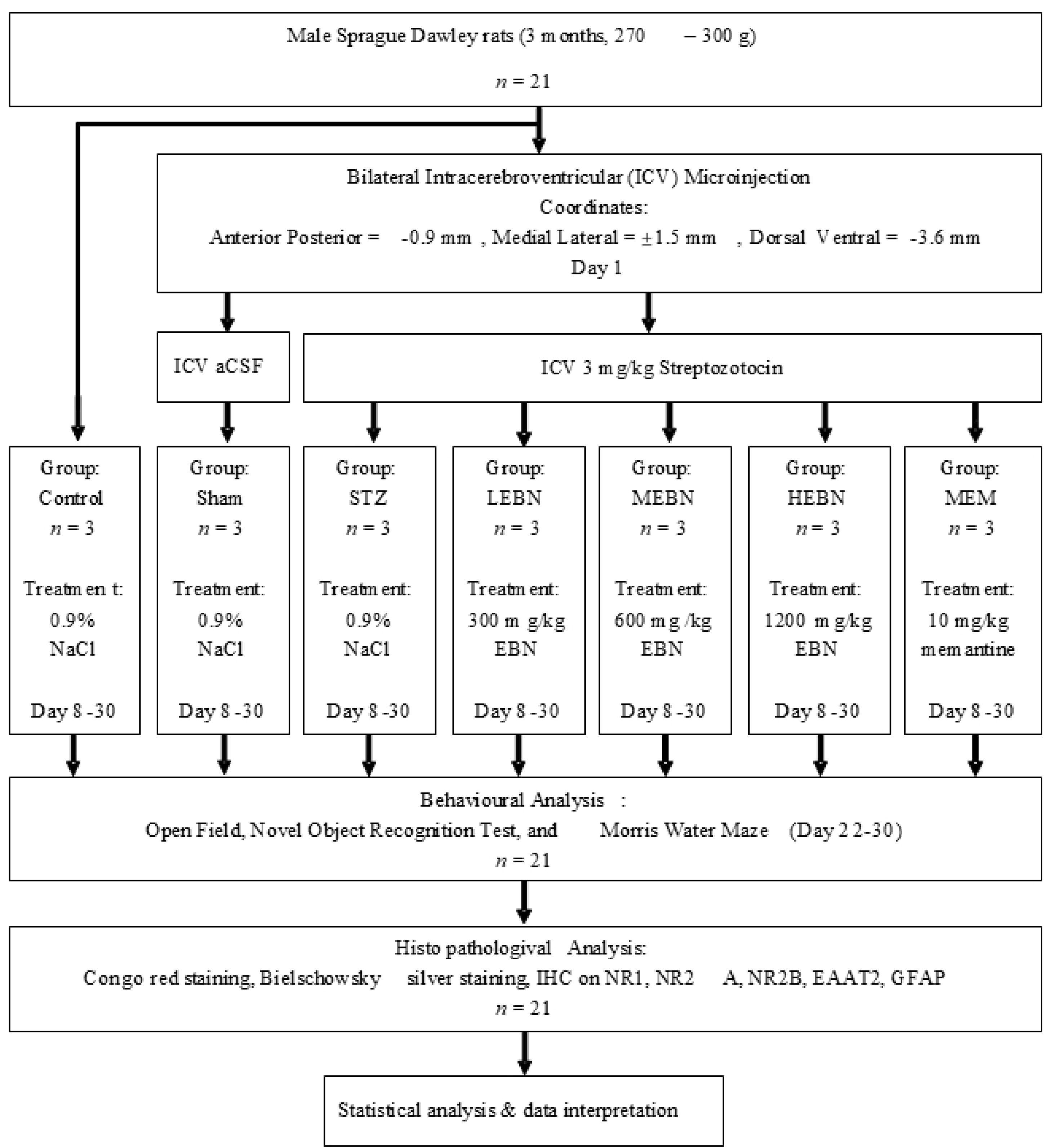
Flowchart of study.

### Validation of Intracerebroventricular Microinjection

Coordinates for the lateral ventricles of postnatal-day 22 (PND 22) rats were initially obtained from an interactive rat brain atlas [19]. Stereotaxic surgery was then performed to validate the coordinates by bilaterally injecting Cresyl violet into the lateral ventricles. Following the procedure, the rat was anesthetized with diethyl ether and decapitated. The brain was harvested, fixed in 10% formalin, and embedded in paraffin. Sagittal sections (3 μm) were prepared using a microtome and examined under an Olympus BX53 microscope.

### Post-Operative Procedure

During the 7-day recovery period, rats were monitored daily for clinical and behavioral signs of pain and infection. To minimize physical trauma following head surgery, food was placed inside the cage for the first 2–3 days. Acetaminophen (100 mg/kg) and tramadol was administered only when necessary [20].

### Behavioural Studies

Throughout the experiments, the observer was blinded to the animals’ group assignments to minimize bias. Illumination was maintained at approximately 10 lux to reduce environmental influence on behavior [21]. A Conexant Video Camera (version 9.071001) was positioned perpendicular to the center of the room to record behavioral activities, which were then analyzed using SMART software (version 3.0.06, Panlab).

### Open Field (OF)

The Open Field Test was conducted following the protocol by Arshad et al. [22] to assess spontaneous motor activity, exploratory behavior, and anxiety-like responses in an open space. The testing room was dimly lit using a fluorescent lamp placed in a corner to minimize floor reflections that could interfere with movement tracking. Rats were habituated to the room for 1 hour before testing. Each rat was placed in the center of an opaque black box (36 × 50 × 36 cm) and observed for 10 minutes. Behavior was recorded using a ceiling-mounted video camera. General activity was evaluated based on horizontal movement, vertical rearing, and total distance traveled, which helped classify motor behavior as normal, hypoactive, or hyperactive. Anxiety levels were inferred from thigmotaxis behavior by measuring the time spent in the periphery versus the center zone (defined as a 20 × 20 cm area in the middle of the field). After each session, rats were returned to their cages, fecal boli were removed, and the box was cleaned with 70% alcohol.

### Novel Object Recognition Test (NORT)

On Day 22, the Novel Object Recognition Test (NORT) was conducted in the same arena used for the Open Field Test, with two identical objects, matched in color, texture, and height, fixed to the floor using tape. The objects were placed 10 cm from the wall and 15 cm apart. Rats were allowed to explore and familiarize themselves with the objects for 10 minutes. On Day 23, one of the familiar objects was replaced with a novel object differing in color, texture, and height, while the other remained unchanged. Rats were again allowed a 10-minute exploration period. All sessions were video recorded, and exploratory behavior was defined as the time spent touching, sniffing, or staring at the object from within 2 cm. The discrimination index (DI) was calculated as DI = (Tn − To)/ (Tn + To), where Tn and to represent the time spent exploring the novel and old objects, respectively. Objects were secured to prevent displacement, and the arena and objects were cleaned with 70% ethanol after each trial.

### Morris Water Maze (MWM)

Three weeks post-surgery, on Day 24 (the day after NORT), the MWM test was conducted following the protocol by Prom-in et al. [23] to assess spatial learning and memory. The test used a circular pool (124 cm diameter, 60 cm height) filled with water to a depth of 30 cm (24 ± 1°C), located in a room with consistent spatial cues. A square escape platform (3×3 inches) was positioned in the ES quadrant, initially placed 1 inch above the water surface for the pre-acquisition trial (Day 0), and then submerged 1 inch below the surface during the acquisition trials (Days 1–5). The pool was divided into four quadrants (NE, ES, SW, WN), and four training trials were conducted daily using randomized start positions from each quadrant. A visual cue was fixed on the north wall to aid navigation. Each trial lasted 60 seconds with 10-minute intertrial intervals. Rats that failed to find the platform within 60 seconds were gently guided to it and allowed to remain for 20 seconds. Following each trial, rats were dried and returned to their cages. On Day 6, a 60-second probe trial was conducted without the platform to assess memory retention of the platform’s location. Key parameters analyzed included path length, mean speed, platform entries, escape latency, and time spent in the target quadrant. Behavior was recorded using a Conexant Video Camera (version 9.071001) and analyzed with SMART software (version 3.0.06, Panlab).

### Histopathological Studies

At the end of the Phase 1 behavioural studies, rats were anaesthetized through decapitation using a guillotine. Brains from all 21 rats were collected for histopathological analysis, fixed in 10% formalin, and embedded in paraffin. Sagittal sections of 10 μm thickness were prepared using a microtome for subsequent analysis.

### Congo Red Staining

Congo red staining was performed to detect Aβ deposits in formalin-fixed brain tissues, following a modified protocol based on Afshar et al. [24]. The tissues were deparaffinized and hydrated in distilled water for 3 minutes, then stained in a working alkaline Congo red solution (1% Congo red in distilled water) for 15 minutes. After staining, the tissues were rinsed in running water for 1 minute and counterstained with Modified Mayer’s Hematoxylin solution for 10 dips. The tissues were then rinsed in running water for 1 minute, immersed in a bluing reagent for 1 minute, and rinsed again in running water. Dehydration and clearing were performed using three changes of absolute alcohol and two changes of xylene, each for 3 minutes. Finally, the tissues were mounted using a resinous mounting medium.

### Bielschowsky’s Silver Staining

Brain tissues were stained using Bielschowsky’s silver staining to detect Aβ plaques, nerve fibers, and tangles, based on Hk Elçioğlu [25]. The tissues were deparaffinized in xylene for 5 minutes per change, followed by rinsing in three changes of absolute alcohol for 1 minute each and then rinsing in running water for 2 minutes. The tissues were incubated in preheated 10% silver nitrate solution at 40°C for 15 minutes, followed by a 2-minute rinse in running water. They were then immersed in preheated ammoniacal silver solution for 10 minutes and agitated in developer solution until a golden-brown color was achieved. The developing process was stopped by quickly immersing the tissues in ammonia water. After a final rinse in running water for 2 minutes, the tissues were immersed in 5% sodium thiosulfate solution, followed by another rinse in running water. Dehydration and clearing were done through three changes of absolute alcohol and three changes of xylene, each for 1 minute. Finally, the tissues were mounted using a resinous medium.

### Immunohistochemistry (IHC)

Brain tissues were sectioned to 3 µm and placed on silanized slides, which were incubated overnight on a slide warmer. The tissues were deparaffinized in two changes of xylene and rehydrated through graded alcohol solutions (100%, 80%, and 70%) for 3 minutes each. After rinsing in running water for 3 minutes, the tissues underwent pretreatment in citrate buffer (pH 6.0) at 110°C for 30 minutes and were rinsed with TBS three times (5, 5, and 8 minutes). The tissues were then incubated with hydrogen peroxide (H_2_O_2_) for 10 minutes, rinsed with TBS three times, and blocked with protein block for 10 minutes. After washing with TBS three times, the tissues were incubated with primary antibodies: 1:250 anti-NR1 (ab193310), 1:100 anti-NR2A (ab174646), 1:100 anti-NR2B (ab93610), 1:1000 anti-EAAT2 (ab205248), and 1:300 anti-GFAP (ab33922) for 30 minutes. Following rinsing with TBS three times, tissues for rabbit monoclonal antibodies (anti-EAAT2 and anti-GFAP) were incubated with a mouse specifying reagent for 10 minutes and rinsed with TBS. All tissues were then incubated with secondary antibody (Goat anti-rabbit HRP conjugate) for 30 minutes, rinsed again with TBS, and incubated with DAB solution for 7 minutes. After immersion in running water for 7 minutes, the tissues were counterstained with Hematoxylin for 50 dips, rinsed again in running water for 7 minutes, dehydrated in an oven for 3 hours, and mounted with a resinous medium.

## STATISTICAL ANALYSIS

Data are presented as mean ± SEM, with statistical significance set at p < 0.05. Normality of the data was assessed using the Statistical Package for the Social Sciences (SPSS) version 27. Changes in body weight were analyzed using one-way ANOVA. Behavioral and biochemical parameters were analyzed daily using a one-way analysis of variance (ANOVA), followed by post-hoc Tukey’s test for multiple comparisons.

## Results

### Validation of Injection Site

A 3 µm coronal section at 0.9 mm posterior to bregma was imaged to validate the ICV injection site (Fig 3). Cresyl violet staining confirmed accurate delivery into the lateral ventricles, located 1.5 mm lateral and 3.6 mm deep from the brain surface, as guided by Paxinos and Watson’s PND21 brain atlas [26]. This validation ensures reproducibility of the STZ-induced SAD model by confirming precise needle placement without affecting adjacent brain regions [27]. The bilateral diffusion of dye supports symmetrical neurodegeneration, aligning with previous findings on ICV-STZ-induced disruptions in hippocampal and cortical insulin signaling [28]. Accurate injection is essential to link observed outcomes directly to central STZ effects, minimizing systemic confounders.

**Fig 3.**
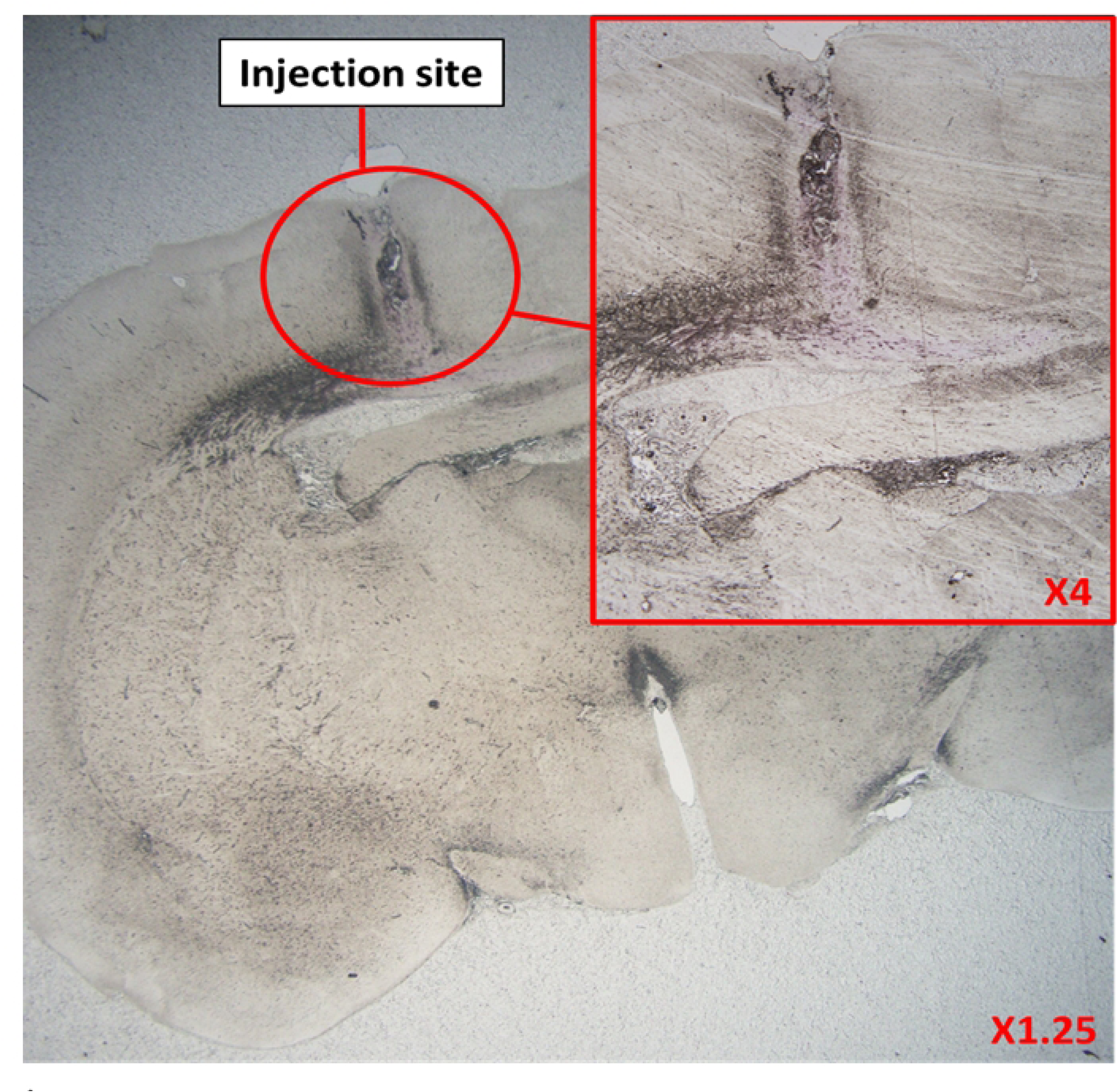
A non-stained, 3 µm coronal section of brain of an ICV rat indicating the intracerebroventricular injection site.

### Assessment of anxiety and locomotion in OF

On Day 22, the open field test assessed locomotion, anxiety, and exploratory behavior (Fig 4). STZ and LEBN groups showed significantly increased path length compared to control, sham, and MEBN, indicating hyperactivity or disorganized exploration. STZ rats also spent more time in the periphery, reflecting heightened anxiety. LEBN and HEBN similarly showed increased thigmotaxis, despite HEBN’s path length being comparable to controls. In contrast, MEBN and MEM normalized locomotor and anxiety-like behaviours, indicating potential anxiolytic and neuroregulatory effects. This supports previous evidence of EBN’s bioactive peptides modulating neurotransmitter systems [29].

**Fig 4.**
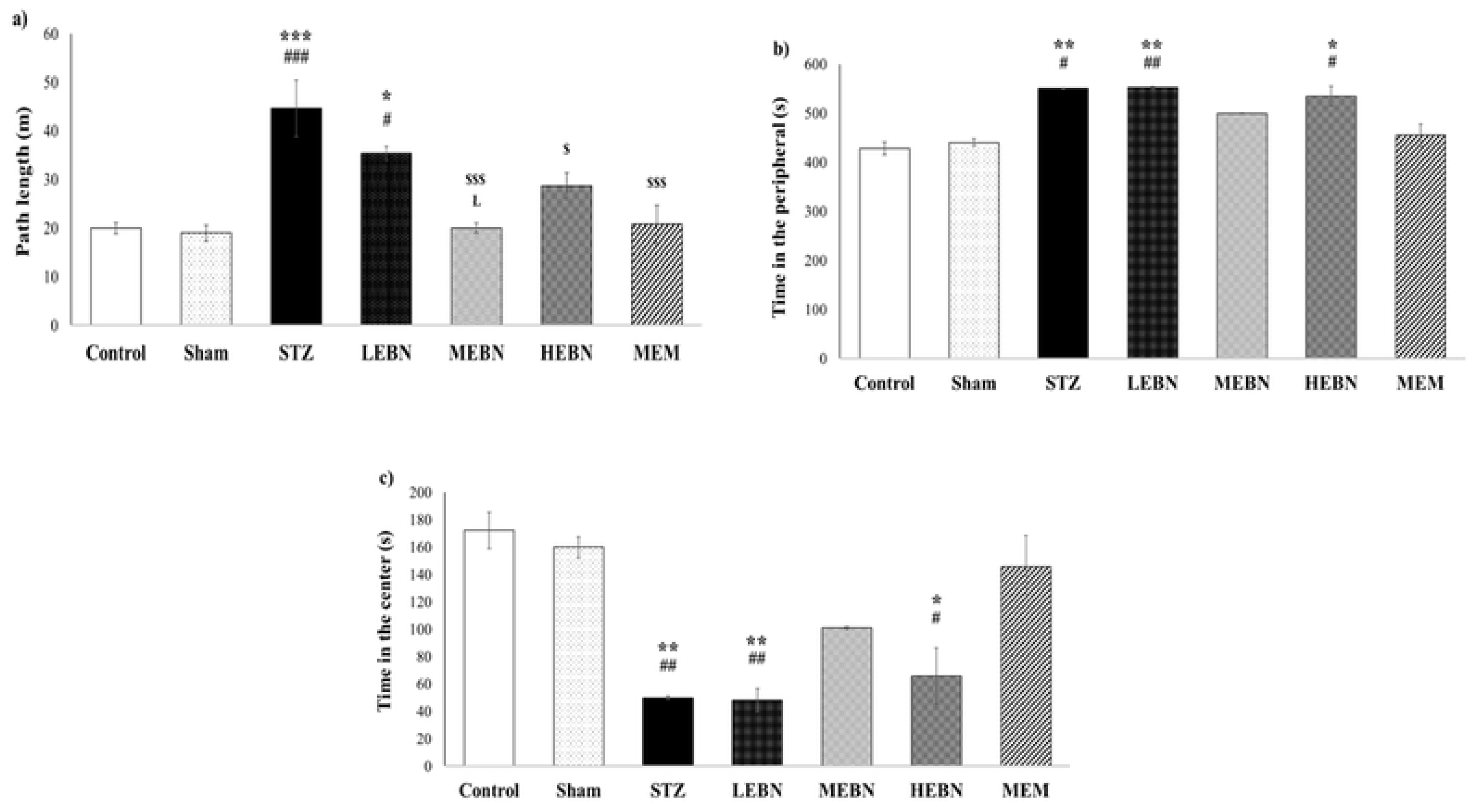
Open field test. a) path length (m), b) time spent in the peripheral (s), and c) time spent in the center (s), where n = 3 per group. Values are expressed as means ± SEM. These parameters were assessed using one-way ANOVA and post hoc Tukey test. *, ** and *** signify a significant difference with p<0.05, p<0.01 and p<0.001 respectively when compared with the Control group. #, ## and ### signify a significant difference with p<0.05, p<0.01 and p<0.001 respectively when compared with the Sham group. $ and $$$ signify a significant difference with p<0.05 and p<0.001 respectively when compared with the STZ group. Ł signifies a significant difference with p<0.05 when compared with the LEBN group.

### Assessment of recognition memory in NORT

In the NORT (Fig 5), STZ and LEBN groups showed significantly higher path lengths than control, sham, MEBN, and MEM, indicating hyperactivity. HEBN had reduced path length compared to STZ. STZ rats spent more time exploring the familiar object (FO), while all EBN-treated groups and MEM showed reduced FO exploration, suggesting partial improvement. STZ, LEBN, and HEBN exhibited impaired novel object (NO) recognition, with significantly lower NO exploration than control and sham. MEBN and MEM significantly improved NO exploration.

**Fig 5.**
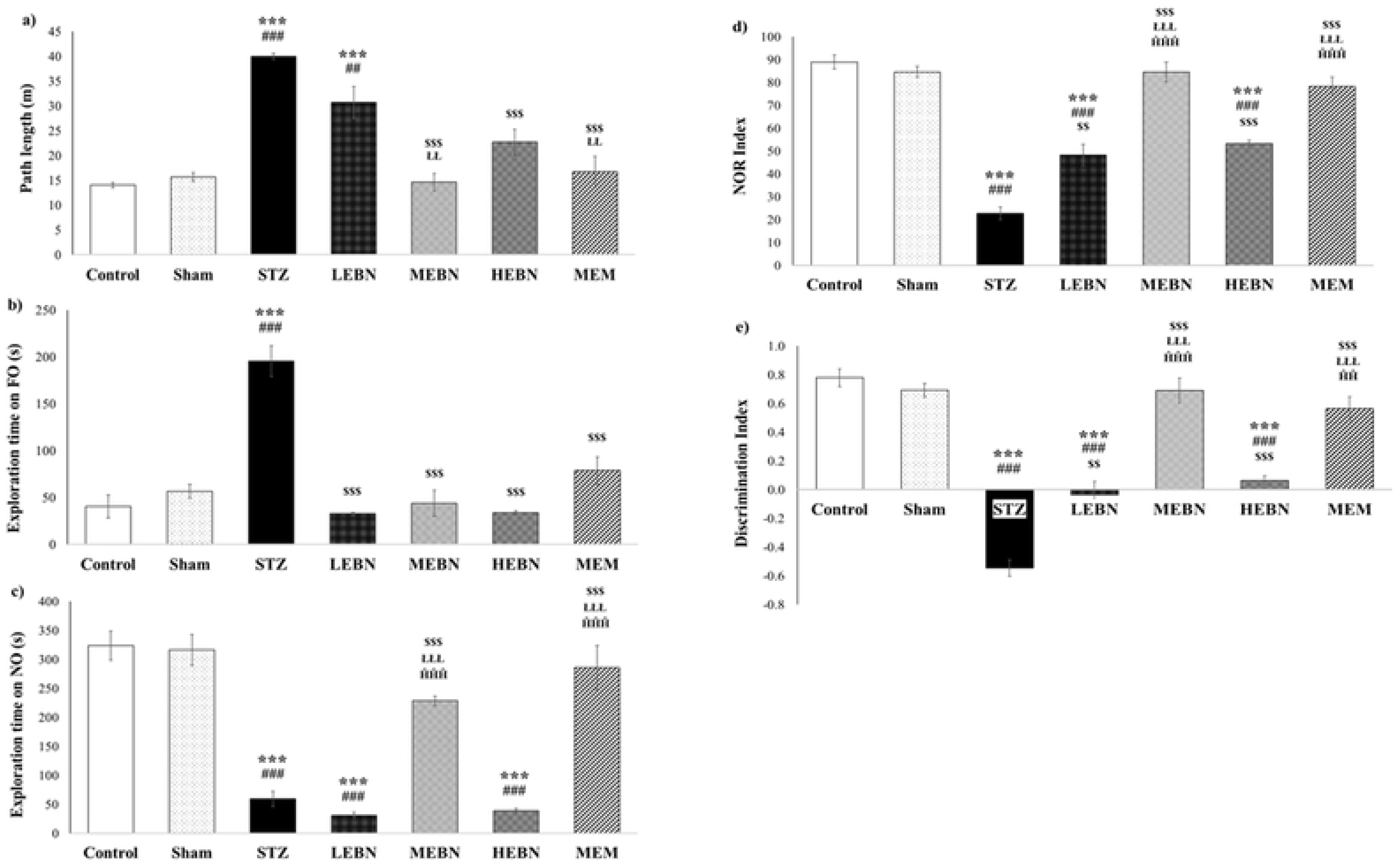
Novel object recognition test. a) Path length (m), b) exploration time on FO (s), c) exploration time on NO (s), d) NOR Index, and e) discrimination index, where n = 3 per group. Values are expressed as means ± SEM. These parameters were assessed using one-way ANOVA and post hoc Tukey test. *** signifies a significant difference with p<0.001 when compared with the Control group. ## and ### signify a significant difference with p<0.01 and p<0.001 respectively when compared with the Sham group. $$ and $$$ signify a significant difference with p<0.01 and p<0.001 respectively when compared with the STZ group. ŁŁ and ŁŁŁ signify a significant difference with p<0.01 and p<0.001 respectively when compared with the LEBN group. ĤĤ and ĤĤĤ signify a significant difference with p<0.01 and p<0.001 respectively when compared with the HEBN group.

The NO and discrimination indices were significantly lower in STZ, LEBN, and HEBN groups, indicating memory impairment. While LEBN and HEBN showed slight improvement over STZ, only MEBN and MEM restored recognition performance to near-control levels. MEBN’s efficacy may be due to optimal receptor modulation and bioavailability, whereas LEBN’s limited effect suggests subtherapeutic dosing [30]. Overall, EBN shows potential in enhancing memory via neurotrophic and anti-inflammatory pathway.

### Assessment of spatial learning and memory in MWM

In the MWM acquisition trial (Fig 6), STZ rats showed significantly longer path lengths, escape latencies, and fewer platform entries compared to control, sham, and most treatment groups, indicating impaired spatial learning. Unlike other groups, STZ rats failed to show improvement across Days 1–5. LEBN rats also exhibited longer escape latencies and reduced platform zone time, particularly on Days 1 and 5, suggesting suboptimal cognitive recovery.

**Fig 6.**
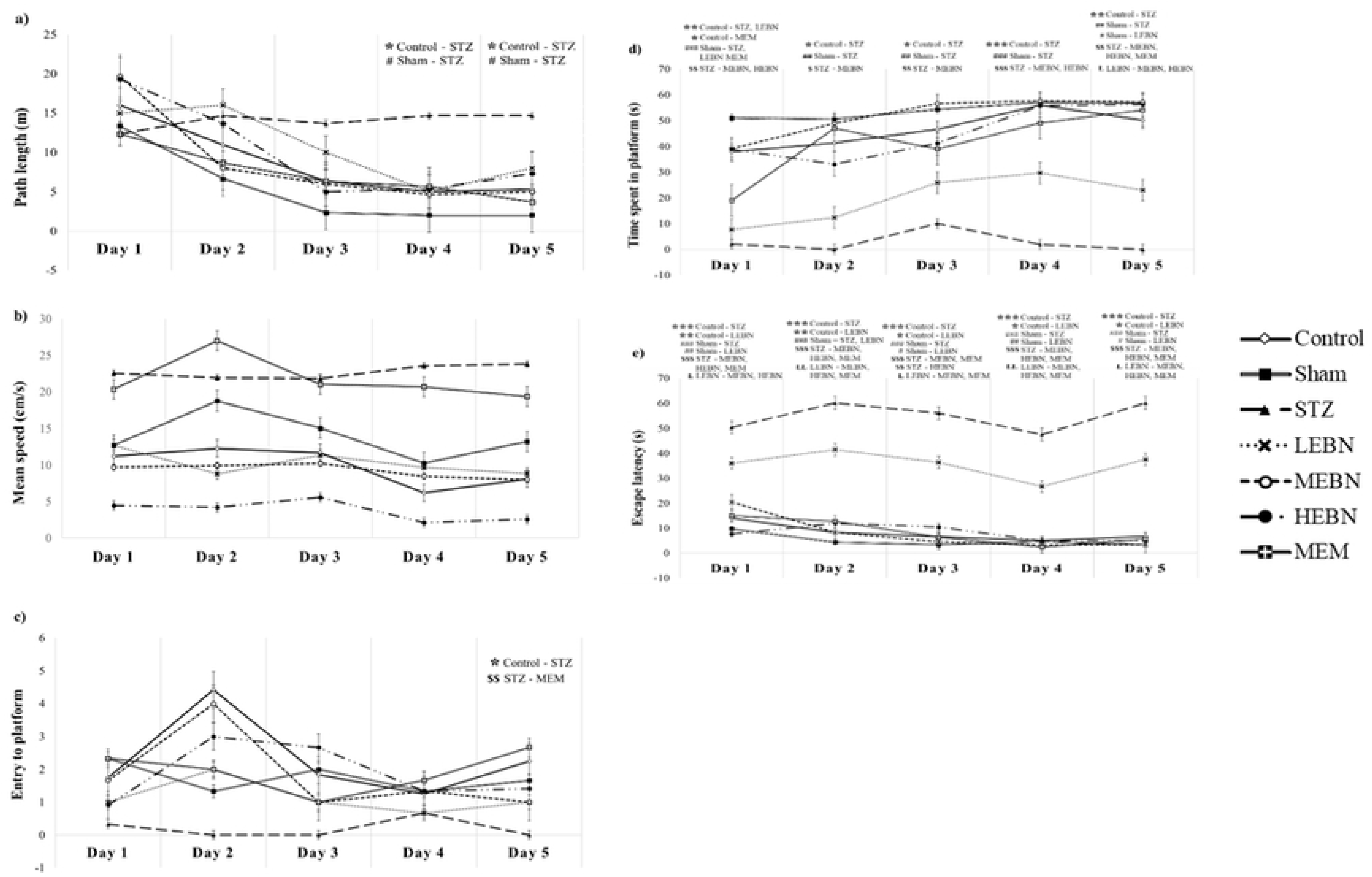
Morris water maze. a) Path length (m), b) mean speed (cm/s), c) entry to platform, d) time spent in platform (s), and e) escape latency (s), during acquisition trial from Day 1 to Day 5, where n = 3 per group. Values are expressed as means ± SEM. These parameters were assessed using one-way ANOVA and post hoc Tukey test. *, ** and *** signify a significant difference with p<0.05, p<0.01 and p<0.001 respectively when compared with the Control group. #, ## and ### signify a significant difference with p<0.05, p<0.01 and p<0.001 respectively when compared with the Sham group. $, $$ and $$$ signify a significant difference with p<0.05, p<0.01 and p<0.001 respectively when compared with the STZ group. Ł and ŁŁ signify a significant difference with p<0.05 and p<0.01 respectively when compared with the LEBN group.

Swimming speed was comparable across groups, except STZ rats swam significantly faster during the probe trial, indicating disoriented, non-goal-directed behavior. During Day 6 probe trial (Fig 7), STZ and LEBN rats spent less time in the platform zone and made fewer platform entries than control, sham, MEBN, and MEM. MEBN showed significantly improved spatial memory, with increased platform entries and reduced escape latency, comparable to MEM.

**Fig 7.**
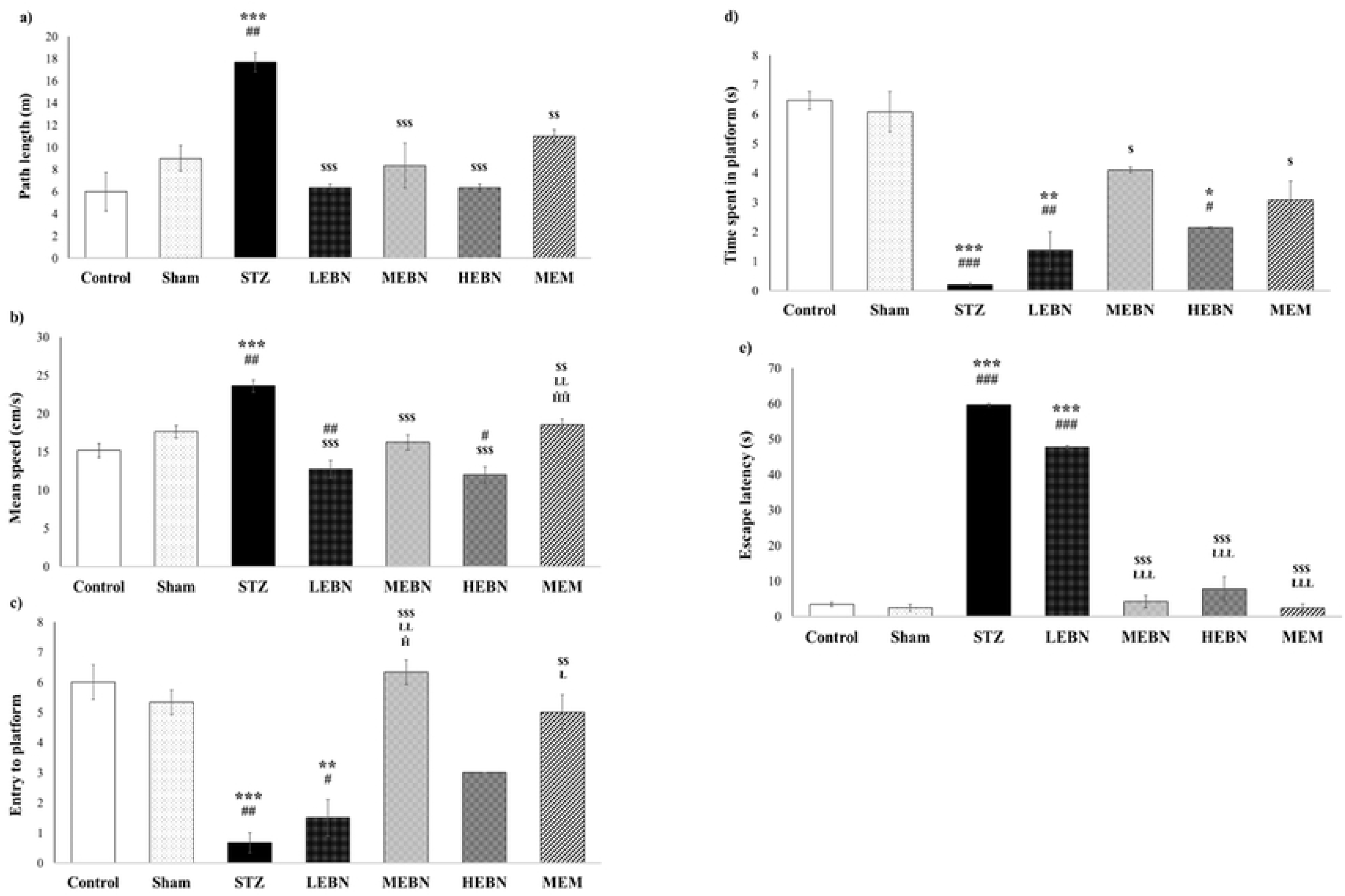
MWM probe trial. a) Path length (m), b) mean speed (cm/s), c) entry to platform, d) time spent in platform (s), and e) escape latency (s), during probe trial on Day 6, where n = 3 per group. Values are expressed as means ± SEM. These parameters were assessed using one-way ANOVA and post hoc Tukey test. *, ** and *** signify a significant difference with p<0.05, p<0.01 and p<0.001 respectively when compared with the Control group. #, ## and ### signify a significant difference with p<0.05, p<0.01 and p<0.001 respectively when compared with the Sham group. $, $$ and $$$ signify a significant difference with p<0.05, p<0.01 and p<0.001 respectively when compared with the STZ group. Ł, ŁŁ and ŁŁŁ signify a significant difference with p<0.05, p<0.01 and p<0.001 respectively when compared with the LEBN group. Ĥ and ĤĤ signify a significant difference with p<0.05 and p<0.01 respectively when compared with the HEBN group.

These findings confirm that ICV-STZ impairs hippocampal-dependent memory [31], while MEBN treatment enhances spatial learning and recall, likely via improved synaptic function and reduced neuroinflammation. LEBN’s limited effect may reflect inadequate CNS penetration or rapid clearance, underscoring the dose-dependent efficacy of EBN.

### Congo red staining

Red-stained deposits (Fig 8), marked by black arrows, were observed in STZ, LEBN, and HEBN rat brains, with STZ showing the highest intensity. These deposits, localized within cell bodies, suggest intracellular Aβ accumulation typical of early STZ-induced AD pathology. MEBN and MEM groups showed minimal staining, comparable to control and sham, indicating reduced Aβ burden. Histopathological analysis using Congo red confirmed Aβ plaque presence, consistent with prior studies [32–34]. However, Congo red results can be inconsistent due to methodological variations and limited clarity under brightfield microscopy [35]. Modified protocols, such as that by Sarkar et al. [36], enhance detection of both plaques and tangles under fluorescence imaging.

**Fig 8.**
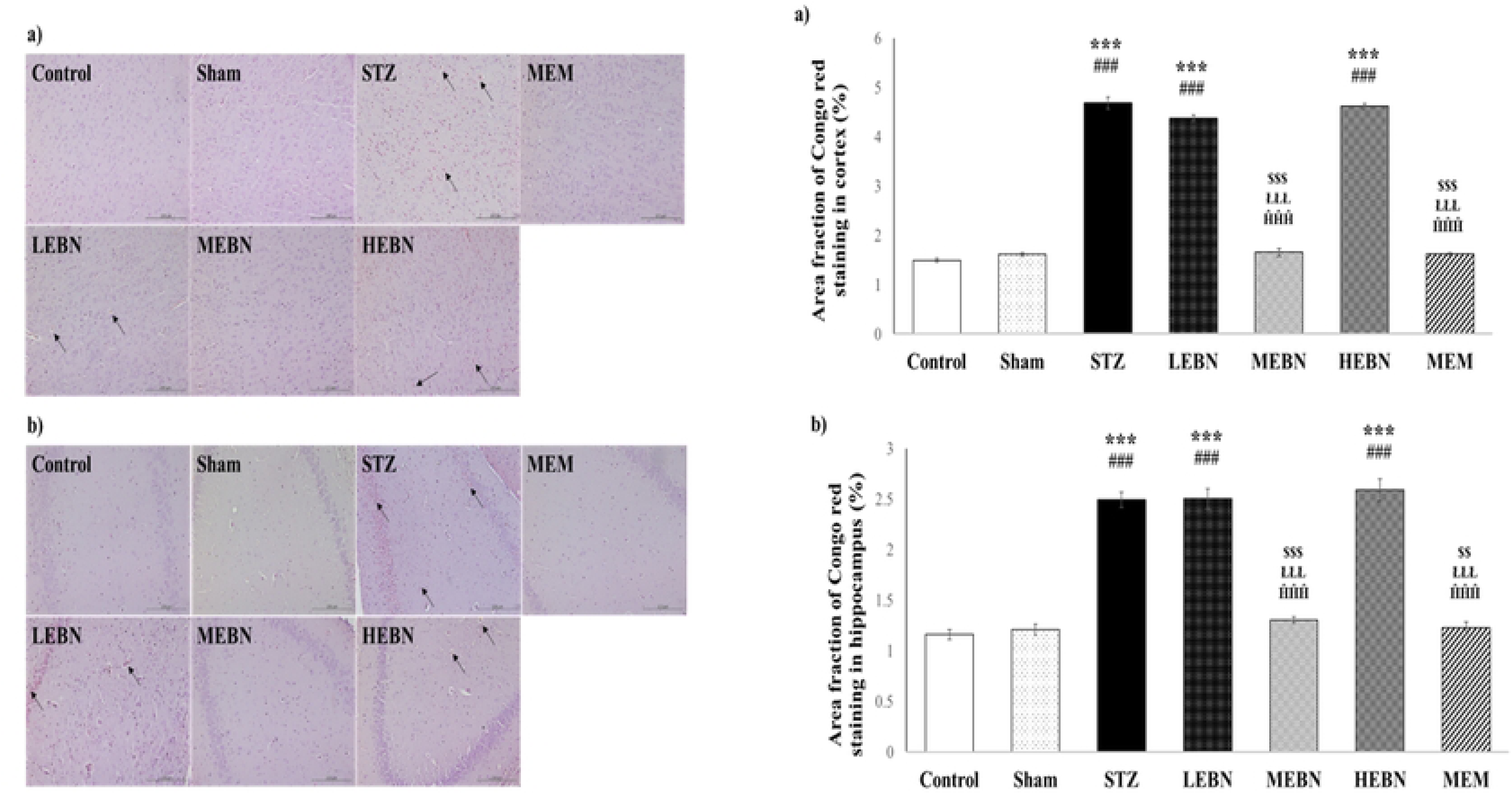
Images show the results of Congo red staining at 20 magnification on rats’ a) cortex, b) hippocampus, where n = 3 per group. Arrow shows amyloid beta plaques stained red. Scale Bar = 200 µm. Values are expressed as means ± SEM. *** signifies a significant difference with p<0.001 when compared with the Control group. ### signifies a significant difference with p<0.001 when compared with the Sham group. $$ and $$$ signify a significant difference with p<0.01 and p<0.001 respectively when compared with the STZ group. ŁŁŁ signifies a significant difference with p<0.001 when compared with the LEBN group. ĤĤĤ signifies a significant difference with p<0.001 when compared with the HEBN group.

### Bielschowsky’s silver staining

Neuritic tangles, indicated by black arrows (Fig 9), were observed in STZ, LEBN, and HEBN groups, with HEBN showing the highest number, followed by STZ and LEBN. These extracellular tangles reflect early-stage neurofibrillary tangle (NFT) accumulation induced by STZ. In contrast, MEBN and MEM groups showed fewer tangles, similar to control and sham, suggesting potential neuroprotective effects.

**Fig 9.**
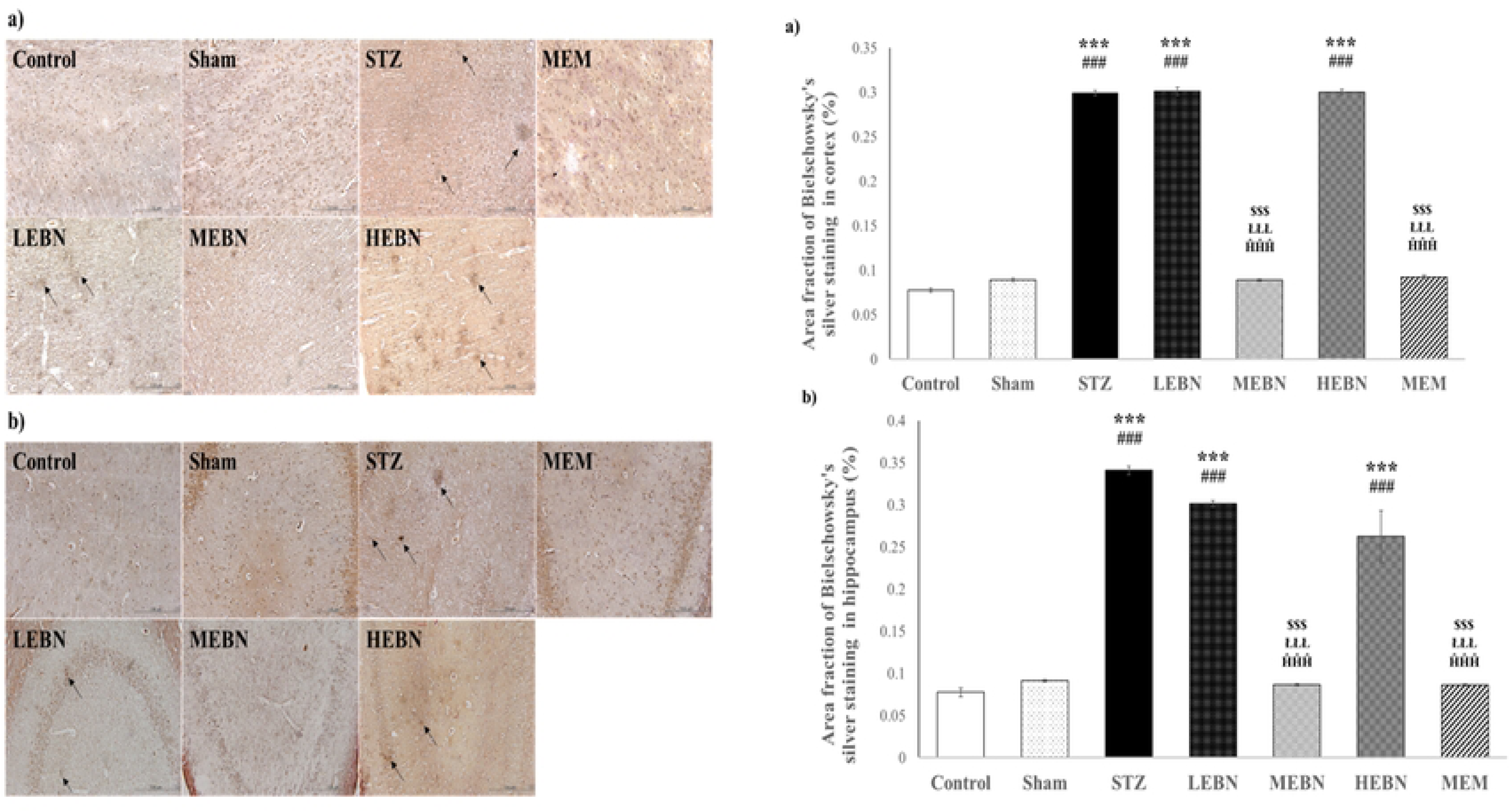
Images show the results of Bielschowsky’s silver staining at 20 magnification on rats’ a) cortex, b) hippocampus, where n = 3 per group. Neuritic tangles showed by the black arrows. Scale Bar = 100 µm. Values are expressed as means ± SEM. *** signifies a significant difference with p<0.001 when compared with the Control group. ### signifies a significant difference with p<0.001 when compared with the Sham group. $$$ signifies a significant difference with p<0.001 when compared with the STZ group. ŁŁŁ signifies a significant difference with p<0.001 when compared with the LEBN group. ĤĤĤ signifies a significant difference with p<0.001 when compared with the HEBN group.

NFT formation was further assessed using Bielschowsky’s silver staining, which revealed argyrophilic spots with stronger intensity in the STZ group, indicating early NFT development [37]. Given the short experimental duration, only pretangles—characterized by fine granular patterns—were detected [38, 39]. Additionally, neuropil thread-like structures and neuritic plaques composed of dystrophic neurites were noted in STZ, LEBN, and HEBN groups. However, higher-resolution techniques like electron microscopy are needed for definitive NFT characterization.

### Effect of EBN on NMDA receptors (NR1, NR2A, NR2B)

Based on IHC images (Fig 10 and 11) GFAP, EAAT2, NR1, NR2A, and NR2B expressions were upregulated in the STZ, LEBN, and HEBN groups. GFAP was highest in STZ, while LEBN showed the highest EAAT2 and NMDAR subunits (NR1, NR2A, NR2B). MEBN and memantine significantly reduced these expressions, approaching control and sham levels. ICV-STZ induced upregulation of NR1, NR2A, and NR2B. The increased NR2A may reflect a compensatory survival response to oxidative stress. Similarly, Yeung et al. [40] reported elevated NR1 and NR2A in STZ-induced AD brains. Wang et al. [41] found NR2B overexpression contributed to cytotoxicity by downregulating CaMKIIα/BDNF/TrkB signaling and reducing synaptic proteins like SYP and PSD-95. Mishra et al. [42] also reported STZ-induced NR1 and NR2B upregulation, which memantine effectively attenuated. GFAP expression was elevated in STZ-treated brains, aligning with increased NFTs and Aβ plaques, consistent with prior studies [42–44]. EAAT2 was highly expressed in STZ rats, with MEBN and memantine normalizing its levels. EAAT2, an astrocytic glutamate transporter, is essential for glutamate clearance (∼95%). Impaired EAAT2 function causes glutamate buildup in the synaptic cleft, overstimulating NMDARs and triggering Na⁺/Ca²⁺ imbalance, excitotoxicity, and neuronal apoptosis [45, 46].

**Fig 10.**
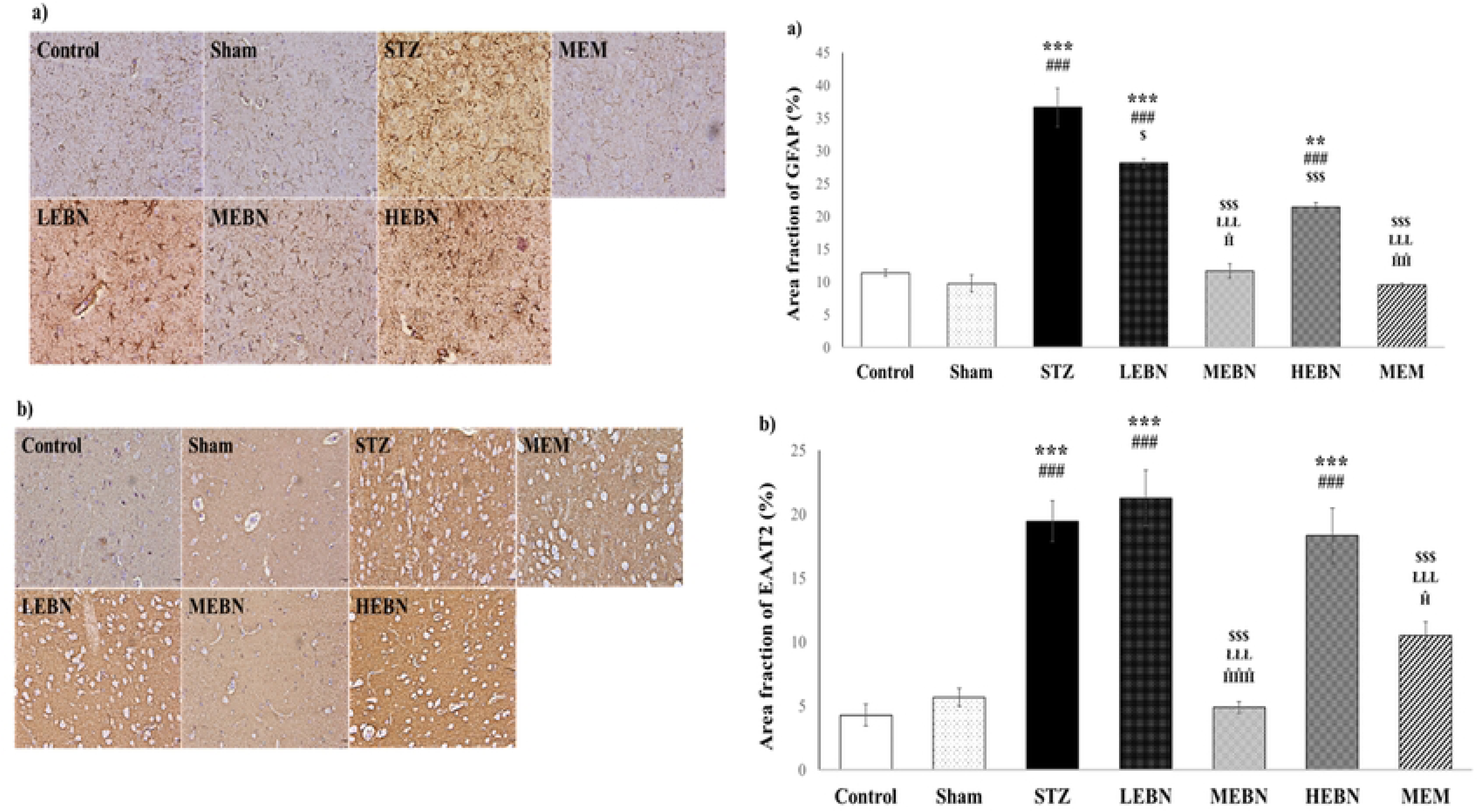
Images show the results of IHC staining for a) GFAP, b) EAAT2, at 20 magnification on rats’ hippocampus, where n = 3 per group. Scale Bar = 100 µm. Values are expressed as means ± SEM. These parameters were assessed using one-way ANOVA and post hoc Tukey test. *, ** and *** signify a significant difference with p<0.05, p<0.01 and p<0.001 respectively when compared with the Control group. #, ## and ### signify a significant difference with p<0.05, p<0.01 and p<0.001 respectively when compared with the Sham group. $, $$ and $$$ signify a significant difference with p<0.05, p<0.01 and p<0.001 respectively when compared with the STZ group. Ł, ŁŁ and ŁŁŁ signify a significant difference with p<0.05, p<0.01 and p<0.001 respectively when compared with the LEBN group. Ĥ and ĤĤ signify a significant difference with p<0.05 and p<0.01 respectively when compared with the HEBN group.

**Fig 11.**
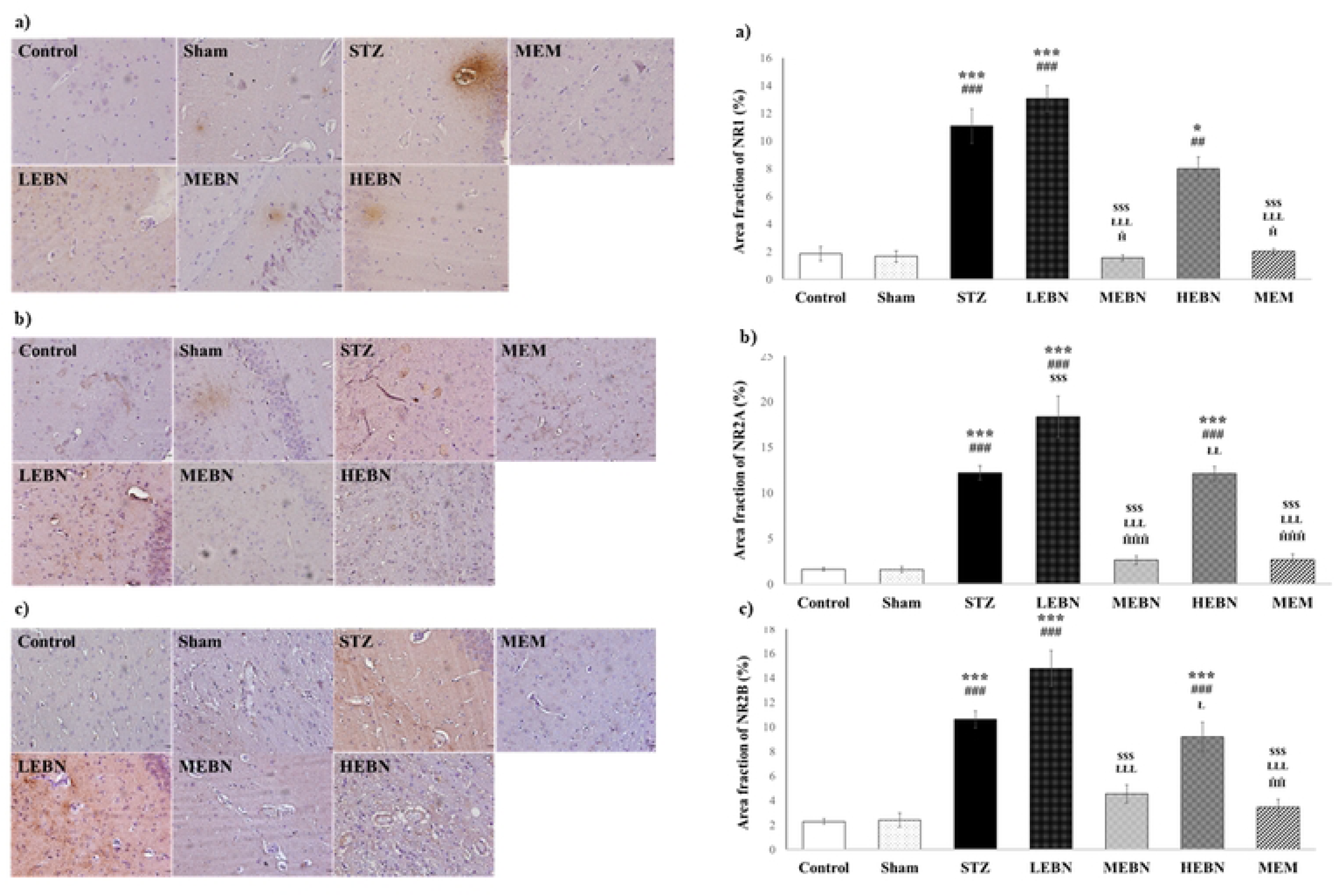
Images show the results of IHC staining for a) NR1, b) NR2A, c) NR2B, at 20 magnification on rats’ hippocampus, where n = 3 per group. Scale Bar = 100 µm. Values are expressed as means ± SEM. These parameters were assessed using one-way ANOVA and post hoc Tukey test. *, ** and *** signify a significant difference with p<0.05, p<0.01 and p<0.001 respectively when compared with the Control group. #, ## and ### signify a significant difference with p<0.05, p<0.01 and p<0.001 respectively when compared with the Sham group. $, $$ and $$$ signify a significant difference with p<0.05, p<0.01 and p<0.001 respectively when compared with the STZ group. Ł, ŁŁ and ŁŁŁ signify a significant difference with p<0.05, p<0.01 and p<0.001 respectively when compared with the LEBN group. Ĥ and ĤĤ signify a significant difference with p<0.05 and p<0.01 respectively when compared with the HEBN group.

## Discussion

The STZ model was selected for its close resemblance to the multifactorial pathophysiology of SAD. A single sub diabetogenic dose of STZ (3 mg/kg) was administered intracerebroventricularly (ICV), inducing brain-specific insulin resistance and SAD-like features in a dose-dependent manner [47]. ICV administration was preferred over intraperitoneal or intravenous routes to target the brain directly, as SAD is considered a brain-specific form of diabetes [48]. STZ was not injected directly into the hippocampus to avoid localized damage and allow for widespread brain distribution via cerebrospinal fluid (CSF), mimicking human SAD more accurately [24]. The 3 mg/kg ICV-STZ dose is well-established to induce memory impairment, cholinergic dysfunction, oxidative stress, impaired glucose metabolism, and neurodegeneration within 21 days, paralleling human SAD pathology [49].

Severe hypoglycemia within the first 24 hours post-STZ injection can cause mortality in rats [50]. In this study, a 25% mortality rate was observed, consistent with previous reports ranging from 7.5% to 33% for 3 mg/kg STZ [51, 52]. To mitigate this, glucose solution was administered via oral gavage for the first three days post-operation. Young male Sprague-Dawley rats (3 months old, 250–300 g) were used due to their sensitivity to STZ [53]. Males were chosen to avoid the neuroprotective effects of ovarian hormones [54].

The hippocampus exhibits functional specialization along its dorsoventral axis: the dorsal hippocampus (dHC) is involved in memory, while the ventral hippocampus (vHC) regulates emotions, including anxiety. The vHC contains glutamatergic, GABAergic, and cholinergic neurons [55], and inhibition of CaMKII-expressing glutamatergic neurons in the vHC has been shown to reduce anxiety-like behavior [56]. Stress activates the hypothalamic-pituitary-adrenocortical (HPA) axis via corticotropin-releasing factor (CRF), oxytocin, and arginine vasopressin (AVP) released from the paraventricular nucleus (PVN) of the hypothalamus [57]. CRF stimulates adrenocorticotropic hormone release from the anterior pituitary, triggering glucocorticoid secretion from the adrenal cortex. Glucocorticoids then exert negative feedback on the HPA axis through receptors in the hippocampus, cortex, and hypothalamus. In the hippocampus, glucocorticoid binding to GR and MR in the ventral subiculum suppresses PVN activity. Accumulation of Aβ plaques and neurofibrillary tangles (NFTs) in the hippocampus can lead to atrophy, elevated cortisol, reduced CRF-expressing cells, and increased CRF1 receptor expression [58].

GFAP is a key astrocytic marker, and its upregulation is a hallmark of reactive astrogliosis. In AD), GFAP levels are elevated in response to the accumulation of amyloid-β (Aβ) plaques and neurofibrillary tangles (NFTs), with increased expression also evident during the early stages of AD, including mild cognitive impairment [43, 59]. Similar patterns of GFAP upregulation have been observed in streptozotocin (STZ)-induced AD models, indicating astrocyte activation as a common feature of AD pathology [28]. One of the critical functions of astrocytes is the regulation of synaptic glutamate levels via EAAT2, the primary glutamate transporter in astrocytes. In AD, impaired EAAT2 function has been reported, resulting in excess synaptic glutamate, which over activates NMDA receptors (NMDARs), disrupts ionic homeostasis, and promotes neuronal apoptosis [45, 46]. Although most evidence points to a downregulation or dysfunction of EAAT2 in AD [60, 61], some studies have reported increased EAAT2 expression in specific contexts [45, 62], possibly reflecting a compensatory astrocytic response. In the present study, we observed elevated expression of NMDAR subunits NR1, NR2A, and NR2B in the hippocampus of STZ-treated rats. This may be linked to axonal transport disruption caused by NFTs, which contributes to neuronal dysfunction and increased vulnerability to excitotoxicity. The resulting neuronal stress could alter glutamate receptor expression as indicated by elevated NR1, NR2A, and NR2B subunits of NMDARs, sensitizing neurons to glutamate-induced damage. Concurrently, the increased EAAT2 expression detected may represent a reactive astrocytic attempt to counterbalance elevated extracellular glutamate levels arising from STZ-induced neuronal injury.

In the present study, at low dose, EBN seems unable to mitigate the neurotoxic effects of STZ. The limited effects of the low dose may be due to its suboptimal bioavailability, as bioactive peptides and glycoproteins in EBN are subject to gastrointestinal degradation, and sialic acid transport across the BBB is receptor-mediated and dose-dependent [63]. Unlike the low dose, there was a significant improvement seen in behavioural, Congo red, Bielchowsky stain, and immunohistochemistry results following MEBN treatment, indicating its clear neuroprotective effect. Given EBN’s multi-medicinal properties, various mechanisms could have been involved in its observed neuroprotective effects. Reduced GFAP seen could be due to its anti-inflammatory effects, by potentially inhibiting the pro-inflammatory cytokine production such as TNF-α [64], which could reduce the glial activation and the overall inflammatory burden in the hippocampus, mitigating the STZ-induced damage. EBN is rich in antioxidants, including lactoferrin, sialic acid, and ovotransferrin [65, 66]. These components can scavenge free radicals and reduce oxidative stress, a key factor in STZ neurotoxicity, which would contribute to improved neuronal function and survival. Through components such as sialic acid, EBN may promote neuronal survival and growth by increasing the expression of neurotrophic factors [67]. Some studies reported that EBN can elevate the expression of Sirtuin-1 (SIRT1) [68], a protein involved in regulating cellular metabolism, stress response, and aging, which has been linked to neuroprotection and improved cognitive function [69, 70]. The reduced Bielchowsky and Congo red staining suggest that mid-dose EBN can mitigate STZ-induced amyloid plaque and NFT formation or promote their clearance, however, this mechanism is not fully understood at present. Moreover, the mid-dose EBN induced decrease in EAAT2, NR1, NR2A, and NR2B expression, might indicate restoration of glutamate homeostasis, rather than a compensatory upregulation. The loss of efficacy at the high dose suggests a potential toxicity threshold or a saturation of beneficial mechanisms. At high concentrations, some antioxidants might become pro-oxidants [71] or interfere with normal cellular signalling pathways [72]. Furthermore, ineffective absorption or metabolism of EBN also could lead to an accumulation of potentially harmful metabolites.

### Conclusion

In conclusion, our study provides evidence for the neuroprotective potential of mid-dose of EBN in the STZ-induced rat model of sporadic AD. The observed significant improvements in behavioural parameters, amyloid plaque and neurofibrillary tangle-like pathology, indicates MEBN’s potential to mitigate key neuropathological hallmarks of SAD. Moreover, MEBN-induced decrease in GFAP suggests a dampening of reactive astrogliosis and concurrent downregulation of NMDARs and EAAT2 compared to STZ group, hints at restoration of glutamate homeostasis. EBN’s antioxidant, anti-inflammatory, and modulation of neurotrophic factors would have most likely contributed to its beneficial effects in this study. The lack of efficacy at low dose and loss of efficacy at high dose of EBN, highlights the importance of dose optimization in harnessing its therapeutic potential. It is important to note that the findings of this study only support the neuroprotective potential of EBN in sporadic AD. To gain a more comprehensive understanding of its potential, future studies should investigate its efficacy in familial AD models and explore the specific bioactive components of EBN responsible for these effects. This would help elucidate the underlying molecular mechanisms and pave the way for potential therapeutic interventions for AD.

### Limitations

The single STZ injection model for AD research has several limitations. Its non-specific toxicity affects both neurons and glia. The acute nature of the model fails to replicate the slow progression of AD. In the present study, the single STZ injection model was chosen to rapidly induce brain insulin resistance and associated early neuropathological changes relevant to AD. Our study aimed to evaluate the potential neuroprotective effects of varying doses of EBN on these early deficits. The consistent poor performance of STZ-treated animals across behavioral assessments (open field, novel object recognition, and Morris water maze) and the significant increases in Congo red and Bielschowsky staining, along with elevated GFAP, EAAT2, and NMDAR subunit expression, confirm the model’s ability to induce AD-related alterations. To address the inherent risks of mechanical damage and potential variability associated with intracerebroventricular injection, a sham-operated control group was included in our study. Importantly, we observed no significant differences in any of the assessed behavioral or biochemical parameters between the untreated control and the sham group. This confirms that the observed deficits in the STZ-treated animals were indeed attributable to the neurotoxic effects of STZ and not a consequence of the surgical procedure itself. The consistently poorer performance of the STZ group compared to the sham group across all measures further validates the model’s utility in inducing AD-related pathology in our experimental setup. Ultimately, the model presents a translational gap due to its poor mimicry of the complex, multifactorial nature of human AD. However, it’s important to note that no current drug-induced model in rats perfectly replicates the full complexity and slow, multifaceted progression of human AD. All of these models often focus on specific aspects of the disease pathology.

## Acknowledgements

The authors would like to thank the members of the Department of Physiology, Department of Pharmacology, and Department of Pathology for their kind assistance.

